# Rapid seasonal evolution in innate immunity of wild *Drosophila melanogaster*

**DOI:** 10.1101/186882

**Authors:** Emily L. Behrman, Virginia M. Howick, Martin Kapun, Fabian Staubach, Alan O. Bergland, Dmitri A. Petrov, Brian P. Lazzaro, Paul S. Schmidt

**Affiliations:** Department of Biology, University of Pennsylvania, 433 S. University Ave. Philadelphia, PA 19104; Janelia Farms Research Campus, Howard Hughes Medical Institute, 19700 Helix Drive Ashburn, VA 20147; Department of Entomology, Cornell University 3125 Comstock Hall Ithaca, NY 14853; Wellcome Trust Sanger Institute, Hinxton, University of Cambridgeshire, CB10 1AS, UK; Department of Ecology and Evolution, University of Lausanne, Lausanne 1015; Department of Biology, Stanford University 371 Serra St, Stanford, CA 94305-5020; Albert-Ludwigs University, Freiburg, Germany; Department of Biology, University of Virginia, 409 McCormic Rd Charlottesville, VA 22904

**Keywords:** Rapid adaptation, innate immunity, *Drosophila melanogaster*, *Thioester-containing protein* 3, *Drosomycin-like 6*, Epistasis, *Providencia rettgeri*, *Enterococcus faecalis*

## Abstract

Understanding the rate of evolutionary change and the genetic architecture that facilitates rapid adaptation is a current challenge in evolutionary biology. Comparative studies show that genes with immune function are among the most rapidly evolving genes in a range of taxa. Here, we use immune defense in natural populations of *D. melanogaster* to understand the rate of evolution in natural populations and the genetics underlying the rapid change. We probed the immune system using the natural pathogens *Enterococcus faecalis* and *Providencia rettgeri* to measure post-infection survival and bacterial load of wild *D. melanogaster* populations collected across seasonal time along a latitudinal transect on the eastern North America (Massachusetts, Pennsylvania, and Virginia). There are pronounced and repeatable changes in the immune response over approximately 10 generations between the spring and fall populations with a significant but less distinct difference among geographic locations. Genes with known immune function are not enriched among alleles that cycle with seasonal time, but the immune function of a subset of seasonally cycling alleles in immune genes was tested using reconstructed outbred populations. We find that flies containing seasonal alleles in *Thioester-containing protein 3 (Tep3)* have different functional responses to infection and that epistatic interactions among seasonal *Tep3* and *Drosomycin-like 6 (Dro6)* alleles produce the immune phenotypes observed in natural populations. This rapid, cyclic response to seasonal environmental pressure broadens our understanding of the complex ecological and genetic interactions determining the evolution of immune defense in natural populations.

## Introduction

The rate at which populations respond to environmental change is a fundamental parameter in the process of adaption. Evolution is historically considered to be an innately slow process that occurs over very long timescales [1], but there are now examples that evolutionary change can occur much faster [2-5]. The limits of how fast populations evolve and the genetic architecture underlying rapid evolution remain unclear [6]. The classical approach to infer adaption through the association of traits and genotypes that co-vary along spatial environmental gradients (e.g., latitude, longitude, altitude) [7] can be expanded across temporal environmental gradients to provide insight to the rate of adaption in the wild.

The biotic environment may shape the rate of adaptation through the immune system, which sits at the crucial interface between an organism’s external and internal environment. Strong selection imposed by pathogens may result in rapid evolution of immune defense in nature because microbiotic infection directly affects host fitness with consequences ranging from resource reallocation away from other functions to host mortality [8-23]. Comparative studies across a broad range of taxa indicate that genes with immune function are among the most rapidly evolving genes in the genome [24-31]. *Drosophila melanogaster* immune genes show evidence of local adaptation across large spatial gradients with high levels of population differentiation and latitudinal enrichment across multiple continents [32-35]. There is less evidence for differentiation at smaller spatial scales [36,37], although some screens of infection response in *D. melanogaster* indicate continental differences in defense quality [36]. Thus, immune defense in natural populations of *D. melanogaster* is a good system to study the how fast natural populations can evolve and genetics underlying the rapid change.

We predict seasonal variation in *D. melanogaster* immune defense even in the absence of established clinal differences in performance. Seasonal climatic changes produce predictable environmental gradients over a temporal scale that select for different phenotypes [38,39] and allele frequencies [40,41] in multivoltine organisms like *D. melanogaster*. Abiotic variables (e.g., temperature) that cycle across seasons can influence microbial growth, so it is possible that microbial communities and pathogen diversity that vary over spatial gradients [42-49] also change as a function of seasonal time [50-53]. Changes in pathogen diversity and frequency across seasons may select for immune resistance or tolerance in either or both of the primary humoral immune pathways: the Toll pathway that is preferentially activated by Gram-positive bacteria or the IMD pathway that is primarily activated by Gram-negative bacteria [54].

We tested whether innate immunity evolves seasonally in mid-Atlantic *D. melanogaster* populations in North America (Massachusetts, Pennsylvania, and Virginia). We found that immune defense changed rapidly and repeatedly from spring to fall, and that seasonally cycling alleles of immune genes determine seasonal variation in resistance to and tolerance of infection. We used reconstructed outbred populations to show that epistatic interactions among seasonally cycling SNPs produced the immune phenotypes observed in natural populations. This rapid, cyclic response to seasonal environmental pressure broadens our understanding of the complex ecological and genetic interactions determining the evolution of immune defense in natural populations.

## Methods

### Experimental Model Details

#### Wild Drosophila Samples

Wild *D. melanogaster* were collected by direct aspiration both in early July (Spring population) and late October (Fall population) at three locations spaced evenly along a 4º latitudinal gradient: George Hill Orchard in Lancaster, MA (42.500493ºN, -71.563580ºE), Linvilla Orchards in Media, PA (39.884179ºN, -75.411227ºE) and Carter Mountain Orchard in Charlottesville, VA (37.991851ºN, -78.471630ºE). Collections were repeated across two years. Isofemale lines were established from wild-caught inseminated females and were maintained on standard cornmeal molasses food under controlled laboratory conditions (25ºC, 12L:12D) on a three-week transfer cycle for 6-8 generations before immune assessment.

#### Recombinant outbred population cages

Recombinant outbred populations [55]fixed for specific seasonal allele combinations in a randomized genetic background were constructed using lines from the Drosophila Genetics Reference Panel (DGRP) [56]. Ten gravid females from 15 lines were pooled to lay eggs for 48 hours for each combination of seasonal alleles. The offspring were permitted to mate freely for at least 10 subsequent non-overlapping generations before immune assessment. This produced populations fixed for the alleles of interest in a heterogeneous unlinked background. The immune function of the two SNPs in *Thioester-containing protein 3 (Tep3)* was tested using three genotypes that combined *2L*:7703202 and *2L*:7705370 (*D. melanogaster* reference genome v.5.39) spring and fall alleles: (1) *Tep3*^*TG*^ contained spring alleles for both *2L*:7703202 and *2L*:7705370, (2) *Tep3*^*TT*^ contained the spring *2L*:7703202 and the fall *2L*:7705370 modifier allele and (3) *Tep3*^*CT*^ contained fall alleles for both SNPs. The final combination of the fall *2L*:7703202 coding allele and the spring *2L*:7705370 modifier allele was too rare in the DGRP to create the recombinant populations. Two independent biological replicate populations were created for each of the three *Tep3* genotypes. Epistatic interactions between *Tep3* and either *Fas-associated death domain (Fadd)* or *Drosomycin-like-6 (Dro6)* were assessed in the same way with recombinant outbred populations fixed for either both spring or both fall *Tep3* alleles and either *Fadd* or *Dro6* alleles.

#### Fly husbandry

Flies were reared in standard laboratory conditions (25ºC, 12:12 L:D) at controlled density in vials. Male flies were collected for infection at 3-5d using light CO_2_ anesthesia. Flies were stored in groups of 10 after infection.

### Method Details

#### Immune survival

Quality of immune defense was probed using systemic bacterial infection [57] with Gram-negative *Providencia rettgeri* [58] and Gram-positive *Enterococcus faecalis* [59] strains that were originally isolated from infected wild-caught *D. melanogaster*. Post-infection survival was measured in males over two repeated blocks of five consecutive days after infection. Mortality was highest in the first 24h and plateaued (Figure S2) so the final mortality 5d post infection was analyzed in the model. Flies were infected with cultures started with a single colony grown to saturation in LB media at 37ºC with shaking overnight and diluted to A_600nm_ of 1.0. Infections were delivered at a dose of 10^3^ to 10^4^ bacteria to each CO_2_-anesthetized fly by inoculating the lateral thorax with a 0.15 mm minute pin (Fine Scientific Tools) dipped into bacterial culture [57]. Two controls were used: a sterile wound by a needle disinfected in 95% ethanol and unwounded flies anesthetized on CO_2_ for the duration of the infection.

#### Bacterial load

The systemic bacterial load of infected flies was quantified using the same infection method as was described above for survival of infection. When evaluating the natural populations, 20 lines from each of the 3 collection locations were infected during a 9a-12p daily infection window. All infections were repeated over two consecutive days by two infectors and the infector and infection order was randomized daily using a random number system. Twelve males from each line were infected each day and maintained in vials with food at 25ºC, 12:12(L:D). The infected flies were measured for bacterial load at 24h after infection. Up to 3 replicate groups of 3 flies were homogenized in 500 mL of LB for the 2012 natural populations and up to three single flies were homogenized in 500 mL of PBS for the 2014 natural and recombinant populations. The samples were then plated on LB agar plates at a dilution of 1:100 for *P. rettgeri*, 1:10 for *E. faecalis* natural populations and 1:1 for the recombinant populations using a Whitley Automatic Spiral Plater (Don Whitley Scientific, Shipley, UK). The plates were incubated overnight at 37ºC and the number of colony forming units on each plate was counted using the ProtoCOL3 automated plate counter (Synbiosis, Cambridge, UK). The number of colonies was used to calculate the concentration of bacteria in each homogenate.

#### Expression data

Expression differences were determined using a published dataset of RNA-seq on 192 inbred sequenced lines from the DGRP [60]. We extracted the expression levels for *Tep3*, *Dro6* and *Fadd* and used the sequence data from [56] to identify the *Tep3*, *Dro6* and *Fadd* haplotypes.

### Quantification and Statistical Analysis

#### Phenotypic statistical analyses

All statistical analyses were performed using the R software (v 3.2.2; The R core team 2012). Post-infection survival was measured daily and the survival 5 days post infection was analyzed using a binomial linear regression. The mean proportion of surviving infected flies was standardized by the survival under sterile wound control treatment and then was evaluated using the following model:

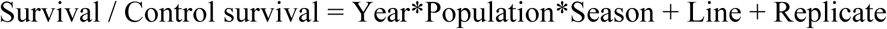

Population, year and season were considered as fixed effects and the random effects of replicate and line were nested within season within population within year.

The number of colonies is used to calculate the concentration of bacteria in each homogenate. The concentrations were log transformed and then analyzed using mixed-model ANOVAs as follows:

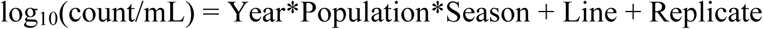

Population, year and season was fixed effects and the random effects were replicate and line nested within season within population within year. Infector and infection order were initially included in the model but had no significant effect and were removed.

#### Seasonal SNPs

Seasonal immune SNPs were identified by screening for alleles that fluctuate in frequency as a function of seasonal time [61] in 88 genes known to have immune function [62]. The seasonal SNPs were cross-referenced with a group of paired spring and fall samples collected from 10 populations along the North American cline by the *Drosophila* Real Time Evolution Consortium (Dros-RTEC 12 unpublished samples; https://sites.sas.upenn.edu/paul-schmidt-lab/pages/opportunities). Additional information was collected on each SNPs including a clinal q-value [61] and a p-value in a genome wide association study to identify SNPs involved with *P. rettgeri* pathogenic infection [63]. Enrichment for immune genes was calculated using customized python scripts that compared proportion of seasonal and non-seasonal immune genes to control genes that were matched for size and position using *χ*^2^ with 10,000 bootstrap iterations.

Linkage disequilibrium (LD) among the candidate seasonal immune SNPs was calculated in the DGRP using allelic correlation of physical distances using the LDheatmap package [64] in *R*. The 205 sequenced inbred lines of the DGRP were used to examine LD among all of the candidate SNPs by chromosome [56].

#### Seasonal genotypes

The genotypes from wild populations were determined using a panel of inbred lines originally collected in Pennsylvania in the spring and autumn of 2012. The lines were inbred by full-sib mating for 20 generations and subsequently sequenced. Genotype deviation was calculated as the difference between observed frequency and a predicted frequency based on the individual alleles. The haplotype distribution of *Tep3* was calculated for SNPs with a minor allele frequency greater than 0.1 using integer joining networks[65] in PopArt vs. 1.7 [66].

#### Expression data

The expression data for *Tep3*, *Dro6* and *Fadd* was extracted from an RNAseq dataset of the DGRP [60]. The lines were sorted by genotype based on the published DGRP data [56] and differences among haplotypes was analyzed using a Welsh t-test in R.

### Results

#### Geographic differences in immunity

The geographic origin of the *D. melanogaster* population across the latitudinal transect determined survival post infection but did not predict systemic bacterial load sustained by flies infected with either pathogen. While survival after *P. rettergi* infection directly depended on the latitude at which the population was collected (*χ*^2^_(2)_=12.805, p=5.87^-4^), geographic origin and season of collection had a combined effect on survival after *E. faecalis* infection (*χ*^2^_(2)_=10.035, p=6.62^-3^). Survival after *E. faecalis* infection was higher in the lower-latitude Virginia population in the spring but the clinal difference disappeared in the fall (Figure 1 A-B). The high-latitude Massachusetts and Pennsylvania populations had similar load and survival after *P. rettgeri* infection and exhibited a greater seasonal change in both survival and bacterial load compared to the lower-latitude Virginia population (Figure 1 C-D).

**Figure 1.**
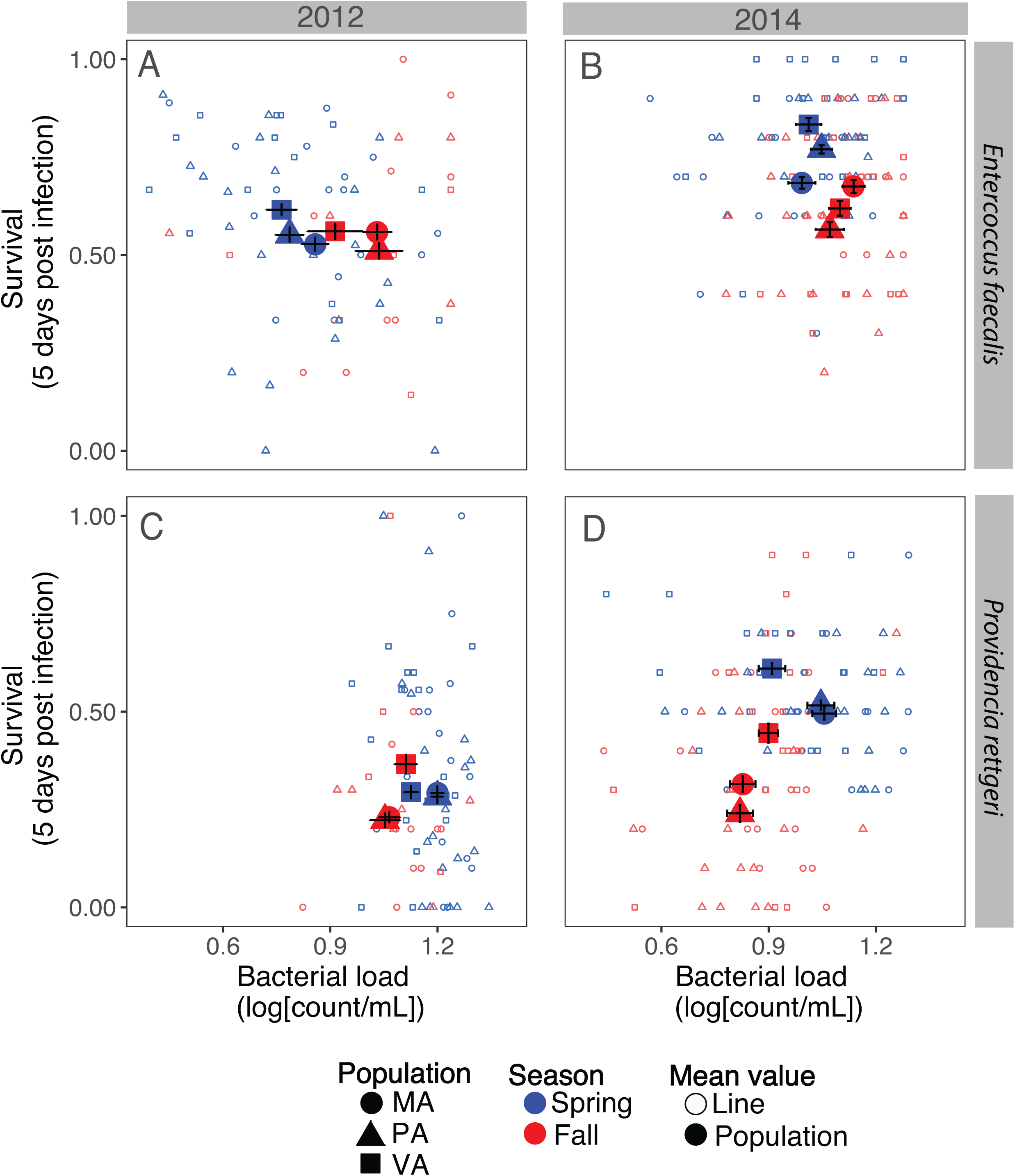
Immune defense relationship between bacterial load and survival in natural spring and fall populations. Isofemale lines (small, outline) were used to calculate population mean (large, filled) from natural orchard populations collected along a latitudinal gradient in Massachusetts (circle) Pennsylvania (triangle) and Virginia (square) in the spring (blue) and fall (red) for two replicate years: 2012 (A & C) and 2014 (B & D). Immune defense was probed with two natural pathogens: a gram-positive bacterium *Enterococcus faecalis* (A&B) and a gram-negative bacterium *Providencia rettgeri* (C&D). Twenty isofemale lines from each collection were measured for 5-day survival after infection and bacterial load at 24 hours post-infection scaled by average load for the experiment.

#### Immunity changes rapidly within a population over seasonal time

Immune defense changed rapidly across approximately 10 generations in the wild from spring to fall. The relationship between bacterial load and survival varied between source population and seasonal collection in a pathogen-specific way (Figure 1). Spring populations were more resistant to *E. faecalis* bacterial growth (F_(1, 219)_=87.758, p<0.0001) and maintained low load with marginally higher survival rates (*χ*^2^_(1)_=3.201, p=07.36^-2^), while the fall populations infected with the same bacteria did not restrict bacterial growth as effectively, resulting in high load and high mortality (Figure 1 A-B). However, the converse relationship occurred when flies were infected with *P. rettgeri*: higher survival in the spring (*χ*^2^_(1)_=16.145, p=5.87^-4^) despite higher bacterial load (F_(1, 215)_=4.3404, p<0.0001) and high mortality in the fall even though the bacterial growth was restricted to low levels (Figure 1 C-D).

#### SNPs in immune genes oscillate across seasonal time

Immune genes as a functional category were not enriched among genes carrying polymorphisms that oscillate in frequency over seasonal time in these populations [61] when compared to controls matched for size and position. We identified 24 candidate SNPs (Table 1) that oscillate in frequency across seasonal time in these populations [61] located within or in proximity to 13 genes that are known to be involved in immune function [67]. Candidate immune genes containing seasonal SNPs were distributed across all levels of the humoral innate immune pathway: two genes in recognition receptors involved with the detection of pathogens, six genes in the signaling cascades and five effector proteins that contribute directly to bacterial killing (Table 1).

**Table 1.** Seasonal immune SNPs identified using whole-genome resequencing of the Pennsylvania spring and autumn populations across three consecutive years. SNPs with a seasonal q-value (SQ) < 0.3 are classified as seasonal and the SNPs investigated here are in bold. Most of seasonal SNPs do not have significant clinal q-values (CQ) and were not significant in a genome wide association study (GWAS) for response to *P. rettgeri* pathogenic infection [52]. The frequency of the SNPs at each collection date is indicated.

| Gene | Position | Effect | Molecular Function | SQ | CQ | GWAS | PA 7.09 | PA 11.09 | PA 7. 10 | PA 11.10 | PA 7.1 | PA 10.11 | PA 11.11 |
| --- | --- | --- | --- | --- | --- | --- | --- | --- | --- | --- | --- | --- | --- |
| Tep2 | 2L:2834400 | Upstream modifier | effector | 0.242 | 0.956 | 0.253 | 0.887 | 0.746 | 0.889 | 0.617 | 0.846 | 0.694 | 0.776 |
| **Tep3** | **2L:7703202** | **NS coding** | **effector** | **0.243** | **0.159** | **0.420** | **0.657** | **0.356** | **0.515** | **0.361** | **0.500** | **0.424** | **0.590** |
| Tep3 | 2L:7703509 | Upstream modifier | effector | 0.151 | 0.529 | 0.084 | 0.840 | 0.567 | 0.813 | 0.667 | 0.838 | 0.677 | 0.694 |
| Tep3 | 2L:7703518 | Upstream modifier | effector | 0.220 | 0.643 | 0.084 | 0.825 | 0.554 | 0.803 | 0.671 | 0.831 | 0.710 | 0.706 |
| Tep3 | 2L:7703748 | Upstream modifier | effector | 0.271 | 0.819 | 0.114 | 0.827 | 0.524 | 0.700 | 0.661 | 0.750 | 0.569 | 0.818 |
| Tep3 | 2L:7703757 | Upstream modifier | effector | 0.291 | 0.956 | 0.632 | 0.748 | 0.476 | 0.488 | 0.541 | 0.664 | 0.367 | 0.618 |
| **Tep3** | **2L:7705370** | **Upstream modifier** | **effector** | **0.219** | **0.163** | **0.385** | **0.479** | **0.158** | **0.255** | **0.240** | **0.457** | **0.273** | **0.444** |
| bsk | 2L:10247834 | Intron | signaling | 0.300 | 0.822 | 0.255 | 0.716 | 0.680 | 0.571 | 0.500 | 0.826 | 0.470 | 0.778 |
| bsk | 2L:10252450 | Intron | signaling | 0.257 | 0.749 | 0.962 | 0.145 | 0.369 | 0.261 | 0.358 | 0.355 | 0.497 | 0.308 |
| Tep1 | 2L:15887030 | Downstream modifier | effector | 0.227 | 0.188 | 0.089 | 0.590 | 0.841 | 0.647 | 0.889 | 0.732 | 0.846 | 0.789 |
| Tep1 | 2L:15888031 | Downstream modifier | effector | 0.221 | 0.520 | NA | 0.000 | 0.368 | 0.063 | 0.360 | 0.000 | 0.013 | 0.012 |
| cact | 2L:16309682 | Downstream modifier | signaling | 0.135 | 0.782 | 0.829 | 0.850 | 0.667 | 0.649 | 0.426 | 0.704 | 0.407 | 0.474 |
| cact | 2L:16310896 | Downstream modifier | signaling | 0.235 | 0.635 | 0.375 | 0.671 | 0.432 | 0.700 | 0.239 | 0.533 | 0.441 | 0.552 |
| cact | 2L:16318067 | Intron | signaling | 0.281 | 0.719 | 0.335 | 0.592 | 0.474 | 0.550 | 0.382 | 0.551 | 0.256 | 0.627 |
| sick | 2L:19923496 | Intron | signaling | 0.232 | 0.032 | 0.505 | 0.096 | 0.047 | 0.130 | 0.048 | 0.328 | 0.053 | 0.269 |
| IM1 | 2R:14270817 | Upstream modifier | effector | 0.256 | 0.695 | 0.423 | 0.358 | 0.075 | 0.571 | 0.193 | 0.213 | 0.115 | 0.390 |
| **Dro6** | **3L:3334769** | **Upstream modifier** | **effector** | **0.201** | **0.427** | **0.000** | **0.770** | **0.613** | **0.814** | **0.612** | **0.798** | **0.489** | **0.625** |
| Drs-1 | 3L:3336529 | Upstream modifier | effector | 0.251 | 0.975 | 0.028 | 0.778 | 0.483 | 0.893 | 0.768 | 0.783 | 0.682 | 0.813 |
| GNBP1 | 3L:18671289 | Downstream modifier | recognition | 0.187 | 0.150 | 0.729 | 0.116 | 0.458 | 0.240 | 0.393 | 0.230 | 0.315 | 0.271 |
| GNBP2 | 3L:18671295 | Downstream modifier | recognition | 0.218 | 0.167 | 0.666 | 0.144 | 0.472 | 0.255 | 0.407 | 0.257 | 0.344 | 0.294 |
| **Fadd** | **3R:17861054** | **UTR 3'modifier** | **signaling** | **0.200** | **0.006** | **0.822** | **0.669** | **0.250** | **0.369** | **0.353** | **0.638** | **0.411** | **0.410** |
| Fadd | 3R:17861073 | UTR 3'modifier | signaling | 0.287 | 0.425 | 0.712 | 0.734 | 0.351 | 0.407 | 0.407 | 0.613 | 0.467 | 0.459 |
| kay | 3R:25600668 | Intron | signaling | 0.200 | 0.588 | 0.743 | 0.686 | 0.453 | 0.607 | 0.464 | 0.636 | 0.383 | 0.475 |
| Tak1 | X:20388404 | Intron | signaling | 0.227 | 0.326 | 0.964 | 0.575 | 0.032 | 0.273 | 0.135 | 0.271 | 0.150 | 0.217 |

#### Seasonally oscillating Tep3 SNPs have functional differences in immunity

Over 1/3 of the seasonally variable SNPs near immune genes were near *Tep* family genes, with *Tep* homologs comprising 1/4 of all of the seasonally variable immune genes. *Tep3* contained numerous seasonally oscillating loci with high LD across the 2.5 kb region in which the seasonal alleles are located in the DGRP (Figure 2B). There were two primary sequence haplotypes carrying spring *Tep3*^*TG*^ variants and two sequence haplotypes carrying the fall *Tep3*^*CT*^ variants in the Pennsylvania orchard (Figure 3F, Table S2). We tested the function of these SNPs using recombinant outbred populations with two loci as markers: the non-synonymous coding change at *2L*:7703202 that is surrounded by five intronic seasonal SNPs and the intronic SNP *2L*:7705370 that is 2 kb downstream from the cluster (*D. melanogaster* reference genome v.5.39). Alleles of the intronic SNP at *2L*: 7703202 were non-randomly distributed with respect to karyotype: in both of the independent DGRP and Pennsylvania populations, we observed that the fall allele (C) was strongly associated with *In(2L)t*. In contrast, the spring allele (T) occurred mostly in a standard arrangement genetic background (Fisher’s exact test; p<0.0001). *2L*:7705730 had no significant association with either arrangement of *In(2L)t* (Fisher’s exact test; p=0.161).

**Figure 2.**
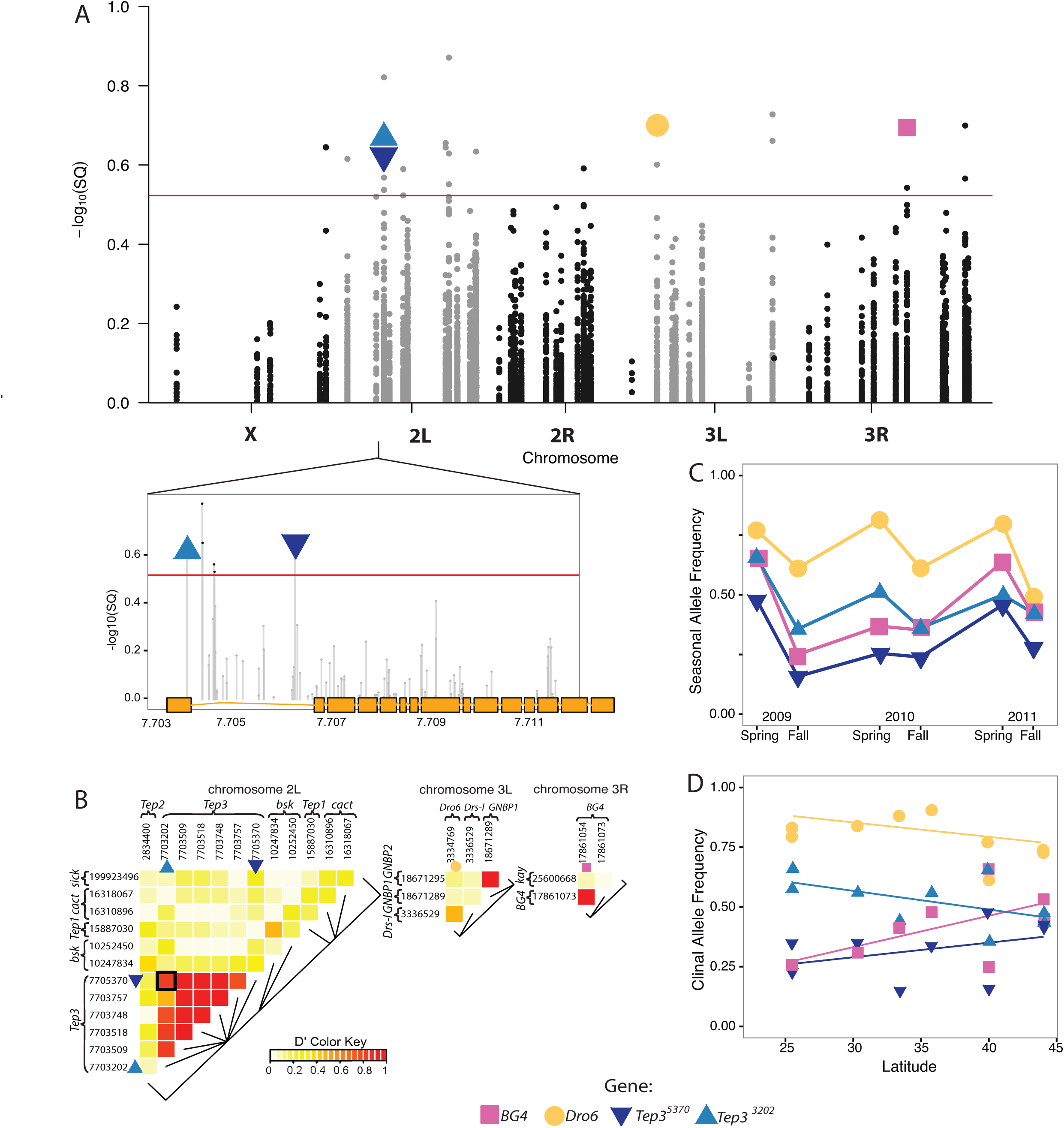
Seasonal changes in immune genes in natural populations. (A) Manhattan plot of SNPs in immune genes that change in frequency as a function of seasonal time with a zoom in on *Tep3*. The red line indicates the seasonal q-value cutoff >0.3[61] and all immune genes that have significant SNPs are labeled by name on the x-axis. The SNPs on which functional analyses were performed are highlighted: *Fadd* (pink square), *Dro6* (yellow circle), *2L*:7703202 (upwards cyan triangle) and *2L*:7705370 (downwards blue triangle). (B) Heat map showing linkage disequilibrium (LD) among SNPs in immune response genes across each chromosome. Linkage disequilibrium calculated as allelic correlation between the physical distances of *2L*:7703202 and *2L*:7705370 in the DGRP is r^2^=0.8138. (C) Cycling of seasonal allele frequencies of candidate immune SNPs across three years. (D) Allele frequencies of candidate SNPs across the latitudinal gradient in the eastern United States. Only *Fadd* shows clinal variation with a clinal q-value of 0.006.

**Figure 3.**
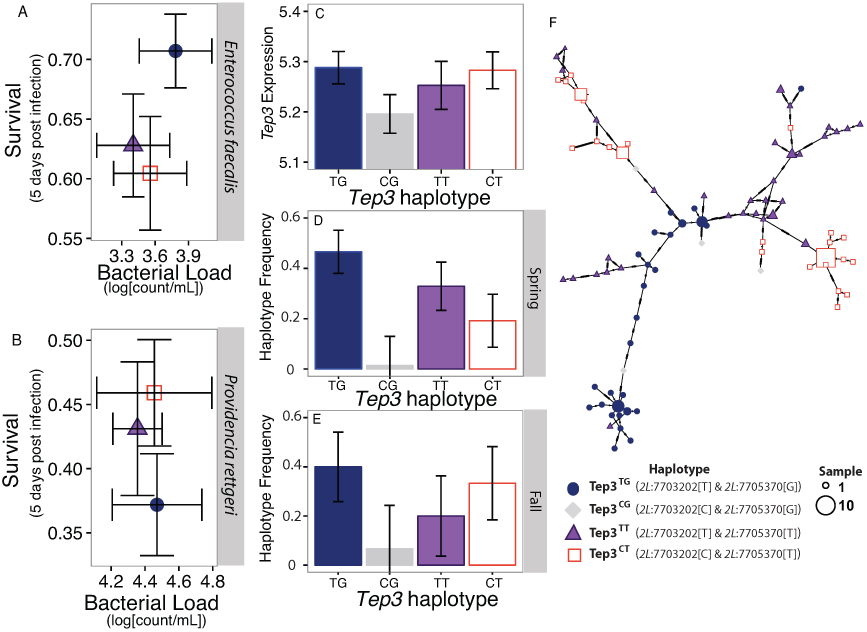
Functional difference of seasonal *Tep3* alleles as defined by the focal SNPs. Mean +/- SE for bacterial load 24 hours post infection and survival 5 days post infection for the *Tep3* genotypes. (A) Higher survival for the spring genotype than the fall or combination genotypes when infected with *E. faecalis.* (B) Additive effect of alleles when infected with *P. rettgeri* (C) Lower constitutive *Tep3* mRNA expression in the rare *Tep3*^CG^ haplotype in flies from the DGRP. (D-E) Frequency of *Tep3* haplotypes in the Pennsylvania orchard across seasonal time. (F) Minimum spanning network illustrates that linkage disequilibrium among the SNPs is maintained in distinct haplotypes.

There was no difference among the *Tep3* recombinant outbred populations in bacterial load, but there was differential survivorship after infection with both Gram-positive and Gram-negative pathogens. Flies containing the spring *Tep3*^*TG*^ haplotype had higher survival than those containing the fall *Tep3*^*CT*^ or mixed *Tep3*^*CG*^ haplotypes when infected with Gram-positive *E. faecalis* (*χ*^2^_(2)_=6.73, p=0.0346; Figure 3A). The *Tep3* SNPs are associated with an additive effect on survival of Gram-negative *P. rettgeri* infection with higher survival in flies containing the fall haplotype than those containing the spring haplotype and intermediate survival in flies containing the mixed haplotype (*χ*^2^_(2)_=3.651, p=0.161, Figure 3B). Flies containing the seasonal *Tep3* haplotypes have no difference in *Tep3* expression in the absence of infection (F_(3, 360)_=1.419 p= 0.239, Figure 3C) based on previously published RNAseq expression of the DGRP lines [60].

#### Epistasis among AMP genes involved in rapid seasonal adaptation

We tested whether additional seasonal SNPs in the immune pathways interact with *Tep3* to facilitate rapid immune evolution across seasons. We examined epistasic interactions in immune function between *Tep3* and a seasonally cycling immune SNP (*3L*:3334769, an upstream modifier of *Drosomycin-like 6* (*Dro6*)), that was shown to significantly affect resistance to *P. rettgeri* in a genome-wide association study [63]. We also tested epistasis among the *Tep3* SNPs and *3R*:17861050, a 3’ UTR modifier in the signaling gene *Fas-associated death domain ortholog* (*Fadd*, also known as *BG4*), which was the only SNP that demonstrated concordant patterns between seasonal change and latitudinal differentiation (Figure 2A, Table 1). There was no difference in immune defense among recombinant populations containing combinations of *Tep3 and Fadd*, but the non-additive interactions among recombinant populations containing *Tep3* and *Dro6* alleles begin to explain more of the complexity of immune defense of natural populations (Figure 4A-D).

**Figure 4.**
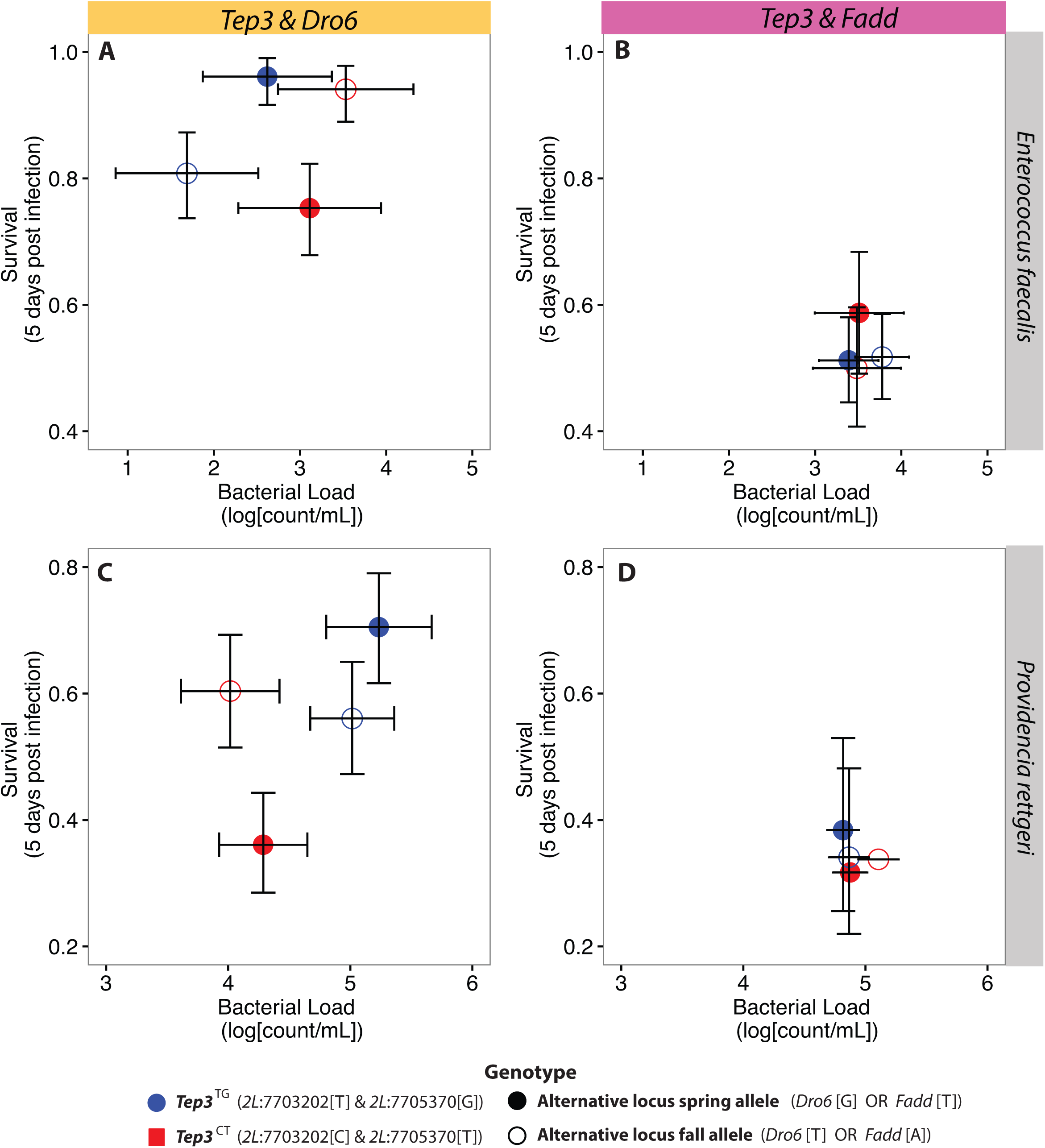
Intergenic interactions among *Tep3, Dro6,* and *Fadd*. Non-additive interaction among *Tep3* and *Dro6* alleles. (A-B). No significant interaction among *Tep3* and *Fadd* SNPs (D-E).

### Discussion

#### Natural populations differ in immunity over geographic space and across seasonal time

We show that immune response differs among populations across space and time. Season of collection is a strong predictor of the immune response across the geographic locations that span 4º latitude with a seasonal decline in resistance to *E. faecalis* and a seasonal decline in tolerance of *P. rettgeri* infection. The change in immunity across seasonal time occurs rapidly within each geographic location with approximately 10 generations between the spring and fall collections. The repeated seasonal change in immune defense is consistent with previous findings for other measurements of stress resistance [38,39]. Together this suggests that the harsh winter selects for a suite of traits that produce a robust spring population and that selection on those traits is relaxed during the summer producing a less stress resistant population in the fall.

Although the strongest differentiation of immunity occurred across seasonal time, there was also a signal of geography along the sampled spatial gradient. Our results contrast with previous studies that did not detect a robust association between latitude and survival [68] or load [36,62]. The difference may be attributed to the interaction between season and latitude. It is possible that geographical differences in immune response may be even greater across a longer distance that may capture a larger difference in pathogen diversity [42-49].

The repeatability of the change in immune defense across replicate years and locations indicate deterministic evolutionary processes. Rearing the lines for multiple generations in a common laboratory environment that is distinct from the external sample sites removes environmental variation and ensures that differences among collections and populations can be attributed to genetic diversity among the source populations. It is possible that gene flow due to migration from other latitudes contributes to the differences between the spring and fall populations. However, migration is unlikely to be the primary cause underlying seasonal immune differences because the latitudinal differentiation was weak compared to seasonal change. Furthermore, infection with different pathogens resulted in opposing clinal patterns but parallel change across seasons. Additionally, migration alone appears insufficient to explain genome-wide differences in allele frequency profiles that characterize spring and fall populations in Pennsylvania orchard [61]; thus, migration is unlikely to explain the seasonal differences in immune response. Wild *Drosophila* populations live in a heterogeneous environment and evolve rapidly in response to environmental parameters that change with season [38,39], potentially including rapid turn-over in microbial and pathogen communities (Figure S2).

#### SNPs in immune genes oscillate across seasonal time

The changes in immune defense are due to differences in genes with immune function across space and time. Genomic screens show that immune genes are enriched across latitudinal gradients [32-35], but we did not find enrichment among immune genes in SNPs that cycle in frequency with season. Seasonal differences in immunity could arise from variation in genes that are not classically identified as part of the immune system and were not detected from our screen. However, the *D. melanogaster* immune system is well characterized and changes in even a single immune gene could affect the phenotypic response to infection even without enrichment for all immune genes. Alternatively, the immune changes may be controlled by non-additive genetic interactions that would not be identified in the enrichment analysis.

#### Immune survival of flies containing seasonally oscillatingTep3 haplotypes

The patterns in the recombinant outbred populations were consistent with the seasonal patterns in natural populations: spring populations and flies containing the spring *Tep3* haplotype both had a higher defense against Gram-positive *E. faecalis* whereas fall populations and flies containing the fall *Tep3* haplotype had higher defense against Gram-negative *P. rettgeri*. Opposite survival patterns for flies with spring and fall *Tep3* haplotypes were consistent with antagonistic pleiotropy [69] within the branches of the immune system limiting the host such that improvements in response to one class of pathogens (e.g., Gram-negative bacteria) restrict the ability to respond to other pathogens (e.g., Gram-positive bacteria). Trade-offs within the immune system occur in several insect systems between humoral antimicrobial peptides that combat microbial infections and phenoloxidase that is deployed against eukaryotic parasites [14,70,71] as well as in the T helper cells of the vertebrate immune system (reviewed in [72]). We hypothesize that genetic variation for allocation of either immune activity may be maintained if the risk of pathogenesis changes over space or time. The genotypes have pathogenic-specific genetic effects. Additivity among the loci in response to *P. rettgeri*, but a non-additive response to *E. faecalis*, suggests that the fall allele at *2L*:7705370, or genetic variants linked to it, has a dominant effect that decreases survival to *E. faecalis* infection.

Our data suggest that these *Tep3* loci are natural variants in immune tolerance because flies containing the haplotypes with the same infection load had differential survivorship. The molecular function of the seasonal loci in *Tep3* remains unclear. *Tep* proteins are α-macroglobulin protease traps that bind to pathogen surface and act as opsonins [73-75]. The polymorphism at *2L*:7703202 produces a nonsynonymous Ala/Val polymorphism at residue 18, but both amino acids produced are hydrophobic. The intronic SNP at *2L*:7705370 is directly upstream of the exon cassette region and may regulate expression, but *Tep3* is constitutively expressed and not strongly induced by *E. faecalis* or *P. rettgeri* infection [76]; B.P. Lazzaro unpublished data). Therefore, the SNPs we examined may most appropriately be considered as markers for a larger haplotype that contains the causal variants.

Pathogen-specific higher survival associated with the spring and fall *Tep3* haplotypes may increase their frequency in the wild compared to flies containing a combination of spring and fall alleles. Inversions could theoretically maintain the LD that preserves the high-fitness spring and fall haplotypes [77,78], but this is unlikely because the *In(2L)t* inversion that contains *Tep3* does not cycle with season [61,79]. Additionally, *Tep3* is not located near a recombination-limiting breakpoint of In(2L)t nor is it in LD with other seasonal immune SNPs within the inversion. However, we found that in two independent populations alleles of the intronic SNP at *2L*: 7703202 were non-randomly distributed with respect to karyotype while *2L*:7705730 had no significant association with either arrangement of *In(2L)t.* LD might be created and maintained by selection against recombinant phenotypes either due to lower immunocompetence or another pleiotropic trait or because of intraspecific genetic incompatibilities. Deleterious incompatibilities maintain distinct haplotypes in *Arabidopsis thaliana* NLR immune receptors [80] and may also explain the near absence of the *Tep3*^*CG*^ combination of spring and fall alleles in all populations examined. Flies containing the *Tep3*^*CG*^ haplotype appear three times across the haplotype tree constructed from the seasonal Pennsylvania inbred lines, suggesting that the haplotype may form occasionally through recombination but does not proliferate in the population. Thus, it is likely that selection for the immune benefits of the spring and fall haplotypes and against the combination of spring and fall alleles maintains these distinct haplotypes in the wild. While these *Tep3* haplotypes explained some of the seasonal differences in immune tolerance of natural populations, other seasonally changing genes may also contribute to the observed differences in bacterial resistance in natural populations

#### Epistasis among AMP genes involved in rapid seasonal adaptation

Intergenic epistatic interactions between *Tep3* and *Dro6* suggest that season-specific genotypes have highest fitness. In our experiment, flies having all spring or all fall alleles had higher survival after infection while flies that contained a combination of spring and fall had higher mortality. This suggests that complex genetic interactions shape winter and summer fitness with distinct haplotypes maintained by non-additive epistatic interactions [81-83].

## Conclusions

With this work, we demonstrate that pathogen-specific innate immunity evolves rapidly in natural populations of *D. melanogaster* across replicate years and geographic locations. Comparative studies across species and among populations have indicated that immune genes evolve faster than other genes in the genome, but the rapid phenotypic and genetic change we observed over approximately 10 generations is a substantially faster rate than previously considered. We tested a small subset of the immune SNPs that oscillate in allele frequency over seasonal time and observed intra- and inter-genic interactions consistent with changes in immune tolerance and resistance across seasons in natural populations, perhaps in response to seasonally changing bacterial communities. Epistatic interactions among seasonally oscillating immune alleles may help facilitate this rapid phenotypic change over a short seasonal timescale. This rapid, cyclic response to biotic variables broadens our understanding of the complex ecological and genetic interactions in the evolutionary dynamics of natural populations.

## Author Contributions

ELB, VMH, BPL & PSS designed the project. ELB & PSS collected the wild samples and ELB & VMH performed the infections. FS analyzed the microbial communities and AOB and DAP inbred and sequenced the seasonal lines used for genotypes in natural populations. ELB, MK and PSS did the data analyses. ELB, VMH, MK, FS, AOB, DAP, BPL and PSS wrote the paper.

## Acknowledgments

This work was supported by NSF GRF DGE-0822 (ELB), the Rosemary Grant Award from the Society for the Study of Evolution (ELB), the Peachey Environmental Fund (ELB), NSF DEB 0921307 (PSS) and NIH R01GM100366 (PSS & DAP).

## Supplemental Material

Figure S1. Related to Figure 1 Post infection survivorship curves for 5 days post infection across seasonal time. Population mean +/- SE.

Figure S2. Related to Figure 1 Microbial community associated with wild and F1 *Drosophila melanogaster* changes over space and time. *D. melanogaster* samples were collected as part of the *Drosophila* Real Time Evolution Consortium (Dros-RTEC 12 unpublished samples; https://sites.sas.upenn.edu/paul-schmidt-lab/pages/opportunities). DNA was extracted as described in [84]. Analysis was performed using a customized MOTHUR (v.1.36.0) [85] script that is available upon request. *Wolbachia* sequences were removed from the analysis.

Table S1. Related to Figure 3. Tep3 haplotypes in the 2012 Pennsylvania population. Focal SNPs are highlighted in black and the genotype combinations are highlighted: spring (blue), fall (red), high-frequency combination (purple), rare combination (grey).

